# The Hippo pathway origin and its oncogenic alteration in evolution

**DOI:** 10.1101/837500

**Authors:** Yuxuan Chen, Han Han, Gayoung Seo, Rebecca Vargas, Bing Yang, Kimberly Chuc, Huabin Zhao, Wenqi Wang

## Abstract

The Hippo pathway is a central regulator of organ size and a key tumor suppressor via coordinating cell proliferation and death. Initially discovered in *Drosophila*, the Hippo pathway has been implicated as an evolutionarily conserved pathway in mammals; however, how this pathway was evolved to be functional from its origin is still largely unknown. In this study, we traced the Hippo pathway in premetazoan species, characterized the intrinsic functions of its ancestor components, and unveiled the evolutionary history of this key signaling pathway from its unicellular origin. In addition, we elucidated the paralogous gene history for the mammalian Hippo pathway components and characterized their cancer-derived somatic mutations from an evolutionary perspective. Taken together, our findings not only traced the conserved function of the Hippo pathway to its unicellular ancestor components, but also provided novel evolutionary insights into the Hippo pathway organization and oncogenic alteration.

## Introduction

A long-standing question in developmental biology is how metazoan species acquired the ability to precisely control their tissue/organ size(Conlon and Raff 1999). This intriguing phenomenon could be traced to some non-metazoan organisms, since their aggregative behaviors can be similarly regulated(Jennings 1940; Olson 2013; Du, et al. 2015). Initially discovered in *Drosophila*, the Hippo pathway is a highly conserved pathway known for its crucial role in organ size control via restricting cell proliferation and inducing apoptosis(Harvey and Tapon 2007; Pan 2010; Halder and Johnson 2011; Yu, et al. 2015; Zheng and Pan 2019), shedding light on this fundamental question in evolution.

In *Drosophila*, the core components of the Hippo pathway comprise two serine/threonine kinases *Hippo* and *Warts*, adaptor proteins *Salvador* and *Mats*, a transcriptional co-activator *Yorkie* and a transcriptional factor *Scalloped*. Together with *Salvador*, *Hippo* phosphorylates and activates *Warts*, which in turn phosphorylates *Yorkie*. *Yorkie* can shuttle between the cytoplasm and nucleus, a process tightly controlled by the *Warts*-mediated phosphorylation. Upon phosphorylation, *Yorkie* is recognized by 14-3-3 proteins and sequestered in the cytoplasm. When Hippo signaling is inactivated, *Yorkie* translocates into the nucleus, where it forms a complex with *Scalloped* to initiate the transcription of genes involved in proliferation and anti-apoptosis, two critical events required for tissue/organ growth. The *Hippo*/*Salvador*-*Warts*/*Mats* kinase cascade can be regulated by multiple upstream regulators. Among them, *Merlin*, *Kibra* and *Fat* have been implicated to play conserved roles in the mammalian Hippo pathway. Moreover, *Hippo* is not absolutely required for the phosphorylation and activation of *Warts*, since another two serine/threonine kinases *Happyhour* and *Misshapen* were also found responsible for *Warts* activation in both *Drosophila* and mammals(Meng, et al. 2015; Tan, et al. 2015; Zheng, et al. 2015), highlighting the complexity of the Hippo pathway regulation.

Although multiple Hippo pathway-related studies have been carried out in species from *Drosophila* to mammals for decades, its components and functions could have deeper conservative roots in evolution. Indeed, several comparative analyses have identified the Hippo pathway components in the ancestor species of cnidarians and bilaterians(Srivastava, et al. 2010; Hilman and Gat 2011; Coste, et al. 2016). In addition, a recent study not only further revealed a non-metazoan origin of the Hippo pathway in evolution, but also uncovered a unicellular organism, *Capsaspora owczarzaki*, where a complete Hippo pathway has already been developed(Sebe-Pedros, et al. 2012). Notably, several Hippo pathway core components including *Mats*, *Hippo* and *Warts* can be even traced to earlier non-metazoan lineages than *Capsaspora owczarzaki*(Sebe-Pedros, et al. 2012). How theses Hippo ancestor components developed their conserved roles in Hippo signaling and how a functional Hippo pathway was evolved from these ancestor components are still largely unknown.

In this study, we traced the evolution of the Hippo pathway organization in both metazoan and non-metazoan species. Specifically, we examined the gene duplication history for the Hippo pathway components from *Drosophila* to human and functionally characterized the unicellular ancestors for the Hippo pathway core components. Given the crucial role of the Hippo pathway in growth control and cancer inhibition, we also systematically analyzed the evolutionarily conserved residues of the Hippo pathway components in human cancer genomes. Taken together, our study not only illustrated the roadmap for the Hippo pathway evolution from its unicellular origin, but also provided insights into the oncogenic alteration of this key tumor suppressive pathway from a novel perspective.

## Results

### Whole genome duplication results in the emergence of paralogous genes for the mammalian Hippo pathway components

Compared to the *Drosophila* Hippo pathway, an interesting phenomenon is the development of paralogous genes for many mammalian Hippo pathway components(Yu, et al. 2015; Zheng and Pan 2019). For example, the *Drosophila* Hippo pathway components *Hippo*, *Warts* and *Mats* were all duplicated into two paralogous genes MST1/MST2, LATS1/LATS2 and MOB1A/MOB1B in mammals, respectively. Some *Drosophila* Hippo pathway components even have more than two paralogous genes in mammals including *Scalloped* (TEAD1/TEAD2/TEAD3/TEAD4 in mammals), *Kibra* (KIBRA or WWC1, WWC2/WWC3 in mammals), *Happyhour* (MAP4K1/MAP4K2/MAP4K3/MAP4K5 in mammals) and *Misshapen* (MAP4K4/MAP4K6/MAP4K7 in mammals). However, as for *Yorkie* (YAP in mammals), *Salvador* (SAV1 in mammals) and *Merlin* (NF2 in mammals), they remained as one gene in mammals.

To elucidate the evolutionary history of the Hippo pathway paralogous genes, we examined the Hippo pathway components in the species ranging from *Drosophila* to human. As shown in Figure 1A, each Hippo pathway component only has one orthologous gene within the tested organisms of Insecta, Branchiopoda, Polychaeta, Clitellata, Gastropoda, Echinoidea and Leptocardii. Interestingly, the emergence of the Hippo pathway paralogous genes was firstly identified in the fish species and they were remained in amphibians, reptiles and mammals (Figure 1A). Among the Hippo pathway components, *Yorkie* and *Merlin* were actually duplicated in fish too, but their paralogs were lost in amphibians and mammals, respectively (Figure 1A); *Taz* was originated in fish and remained as one gene during evolution (Figure 1A); no paralogous genes of *Salvador* were identified in all the tested species from *Drosophila* to human (Figure 1A). Collectively, these data indicate that fish is the turning point for the Hippo pathway in evolution, where many Hippo pathway components acquired their paralogous genes.

**Figure 1.**
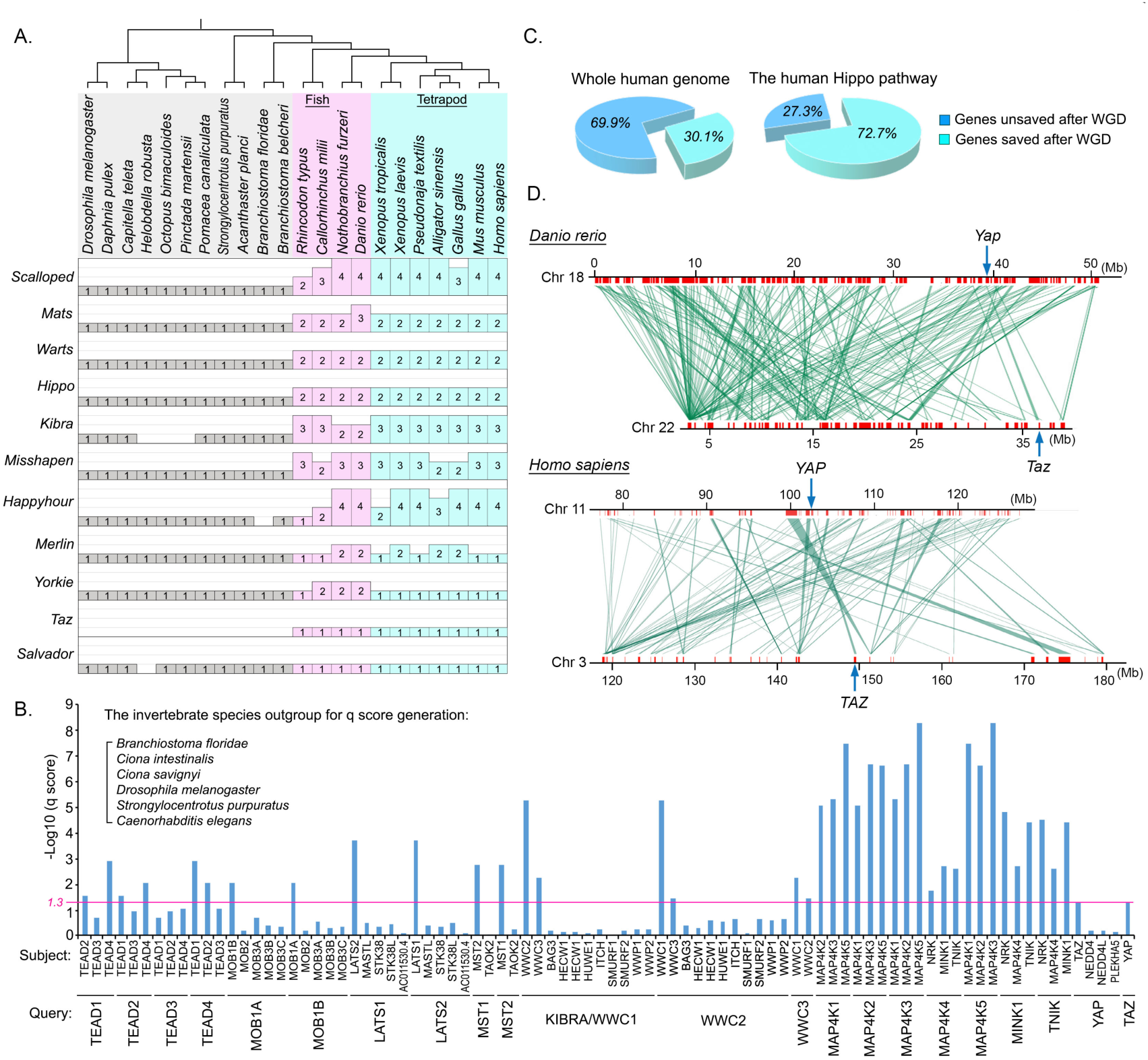
The development of the Hippo pathway paralogous genes is caused by the whole genome duplication in fish. (This figure is related to Figure S1 and Tables S1-S3) **(A)** Illustration of the Hippo pathway paralogous genes in the indicated species from *Drosophila* to human. The Hippo pathway components were searched in the indicated species’ genomes by TBLASTN. The Hippo paralogous gene number in each indicated specie is shown. The listed species in Protostomia, Echinodermata and Leptocardii were labeled in grey; the listed species in Fish were labeled in pink; the listed species in Terapod were labeled in light blue. **(B)** The development of the Hippo pathway functional paralogous genes was a result of whole genome duplication. The q scores among the Hippo pathway functional paralogous genes were more significant than those of control genes. **(C)** The Hippo pathway is enriched with the duplicate genes saved after whole genome duplication as compared with the whole human genome. **(D)** Genomic regions near *YAP* and *TAZ* genes in both *Danio rerio* and *Homo sapiens* show a strong syntenic relationship. The neighboring gene information of *YAP* and *TAZ* were obtained from BioMart.

Notably, the Hippo pathway paralogous gene number in fish species was highly correlated with the rounds of whole genome duplication (WGD) (Figure S1A), including three rounds of WGD happened in different fish ancestors and two in the vertebrate ancestors(Glasauer and Neuhauss 2014). To determine whether WGD resulted in the emergence of the Hippo pathway paralogous genes in human, we analyzed the statistical confidence q-score for each paralog pair of the Hippo pathway genes in human genome, where several invertebrate species were included as an outgroup. As shown in Figure 1B, the q scores generated among the Hippo pathway paralogous genes were more significant than the ones generated between these Hippo pathway genes and control genes, suggesting that these Hippo pathway paralogous genes were emerged as a result of WGD. Moreover, compared with the average rate of genes saved after WGD in the human genome (30.1%)(Tinti, et al. 2014), around 72% of the Hippo pathway components retained their duplicated genes after WGD (Figure 1C). Taken together, these data suggest that the Hippo pathway is highly enriched with paralogous genes after the WGD in vertebrate ancestors, allowing its stable role in regulating tissue/organ homeostasis within the following species such as amphibians, reptiles and mammals in evolution.

### YAP and TAZ share the same evolutionary origin

By analyzing the evolutionary history of the Hippo pathway paralogous genes, we noticed that TAZ was firstly identified in fish (Figure 1A) and it shares a significant q-score with YAP in the WGD gene analysis (Figure 1B). Moreover, YAP and TAZ are similarly regulated and can function redundantly in the mammalian Hippo pathway. These evidences suggest that TAZ could be developed from *Yorkie* during WGD. To test this hypothesis, we performed a syntenic analysis of the surrounding regions of *YAP* and *TAZ* genes in both fish and human genomes, where the ancient vertebrate *Ciona intestinalis* genome was included as an outgroup (Figure S1B). Interestingly, the genes surrounding *Yorkie* in *Ciona intestinalis* genome were similar to the genes near YAP and TAZ in both fish and human genomes (Figure S1B). Moreover, the genomic regions near YAP and TAZ showed a strong syntenic correlation between fish and human genomes (Figure 1D). These data demonstrate that TAZ and YAP are paralogous genes because of WGD.

In addition to the redundant roles between YAP and TAZ in the Hippo pathway, accumulated evidences also indicated that YAP has a stronger influence than TAZ(Plouffe, et al. 2018). To elucidate the functional divergence between YAP and TAZ, we examined their coding sequence divergence by using the program DIVERGE 3.0(Gu, et al. 2013). Interestingly, the YAP and TAZ coding sequences belong to the Type I divergence with the coefficient (θ_1_) significantly greater than 0 (Figure S1C). This finding suggests a potential site-specific rate shift between YAP and TAZ after the gene duplication, which may lead to their partially functional difference.

### Roadmap of the Hippo pathway evolution in unicellular species

Next, we further explored how a complete Hippo pathway was evolved from its ancestor components. Consistent with a previous study(Sebe-Pedros, et al. 2012), most of the Hippo pathway core components can be identified in unicellular species (Figure 2A). Moreover, a functional and complete Hippo pathway has already emerged in an unicellular organism *Capsaspora owczarzaki*(Sebe-Pedros, et al. 2012). Actually, several Hippo ancestor components can be traced to even earlier unicellular species than *Capsaspora owczarzaki* (Figure 2A): *Mats* is the earliest appeared Hippo pathway component that can be identified in the unicellular species such as *Tetrahymena thermophile*, *Thalassiosira pseudonana, Chlamydomonas reinhardtii*, *Trichomonas vaginalis*, *Naegleria gruberi*, *Arabidopsis thaliana*; following that, *Hippo* was the next Hippo pathway component appearing in Amoebozoa (e.g. *Acanthamoeba castellanii*), Diclyostelia (e.g. *Dictyostelium discoideum*) and Apusozoa (e.g. *Thecamonas trahens*); another Hippo pathway kinase cascade component *Warts* was firstly identified in *Fonticula alba* as well as the following fungi species (e.g. *Saccharomyces cerevisiae*), while both of these two species did not contain *Hippo*. After these three Hippo components, the kinase cascade (i.e. *Hippo*/*Warts*/*Mats*) and transcriptional complex (i.e. *Yorkie*/*Scalloped*) in the Hippo pathway became completed in *Spizellomyces punctatus* and *Corallochytrium limacisporum*, respectively. Finally, an intact Hippo pathway was eventually developed in *Capsaspora owczarzaki*. (Figure 2A).

**Figure 2.**
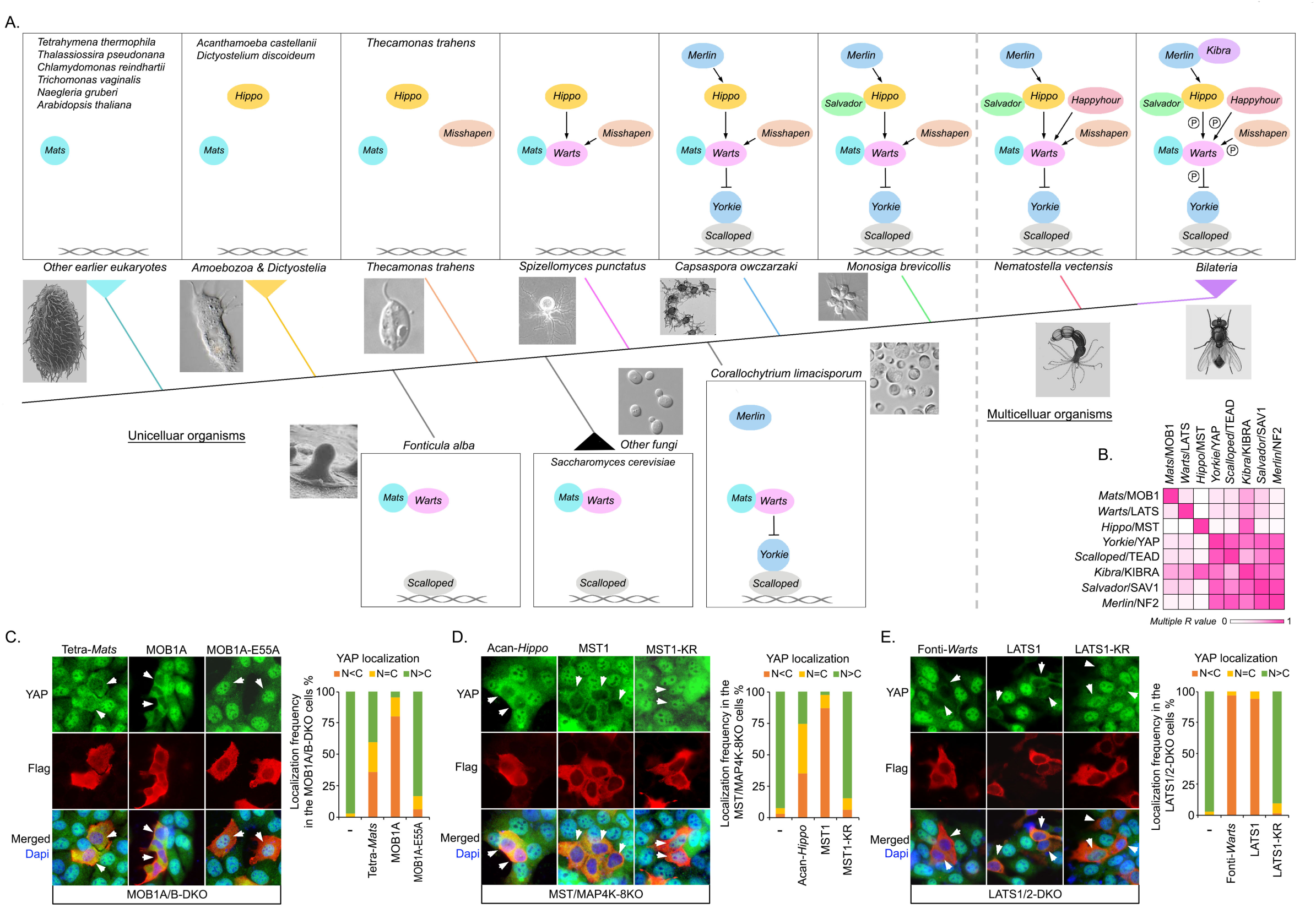
The Hippo pathway unicellular ancestor components show conserved activities in human cells. (This Figure is related to Figures S2 and S3; Tables S1 and S4) **(A)** Schematic illustration of the Hippo pathway evolution from unicellular organisms to Bilateria. The Hippo pathway components were searched from the genomes of the indicated species ranging from *Tetrahymena thermophila* to Bilateria by TBLASTN. The color of lines was matched with the firstly emerged Hippo pathway component in the indicated species. **(B)** Coevolution analysis among the Hippo pathway core components. Linear regression analysis was performed to determine the correlation between pairwise evolutionary distances based on a multiple protein sequence alignment. (**C-E**) Three Hippo pathway ancestor components show conserved functions in the human cells. The Hippo pathway unicellular ancestor genes *Mats* (from *Tetrahymena thermophile*), *Hippo* (from *Acanthamoeba castellanii*) and *Warts* (from *Fonticula alba*) were synthesized and respectively expressed in the MOB1A/B double knockout (DKO) (**C**), MST/MAP4K-8KO (**D**) and LATS1/2 DKO (**E**) HEK293A cells. Immunofluorescence was performed using YAP and Flag antibodies. The human MOB1A, MST1 and LATS1 were taken as positive controls, while their inactive mutant MOB1A-E55A, MST1 kinase-dead mutant (K59R) and LATS1 kinase-dead mutant (K734R) were included as negative controls. Flag-positive cells (arrows) from ~30 different views (~200 cells in total) were randomly selected and quantified for YAP localization.

We also examined the co-evolutionary relationship among the Hippo pathway core components by analyzing their gene sequences from unicellular organisms to human. Interestingly, the earliest emerged Hippo pathway components *Mats*/MOB1, *Warts*/LATS and *Hippo*/MST showed a relatively weak correlation in between and with other Hippo pathway components as indicated by the low multiple R value (Figure 2B). In contrast, strong coevolution was identified among the Hippo nuclear components (*Scalloped*/TEAD and *Yorkie*/YAP), one kinase cascade adaptor (*Salvador*/SAV1) and two upstream regulators (*Merlin*/NF2 and *Kibra*/KIBRA). This result indicated that the intrinsic function of *Mats*/MOB1, *Warts*/LATS and *Hippo*/MST could be established independently at their evolutionary origins, whose upstream and downstream Hippo pathway components were then coevolved, allowing the development of a regulatory network and a functional output for these three Hippo pathway ancestor components.

To test this hypothesis, we characterized the roles of the *Mats*, *Hippo* and *Warts* ancestor proteins in *Tetrahymena thermophile* (hereafter called its *Mats* as “Tetra*-Mats*” in short), *Acanthamoeba castellanii* (hereafter called its *Hippo* as “Acan*-Hippo*” in short) and *Fonticula alba* (hereafter called its *Warts* as “Fonti*-Warts*” in short), respectively, in their corresponding human knockout (KO) cells. Interestingly, both Tetra*-Mats* and Acan-*Hippo* were able to partially rescue YAP’s cytoplasmic localization in the MOB1A/B double KO (DKO) (Figure 2C) and MST/MAP4K-8KO HEK293A cells (Figure 2D), respectively. Surprisingly, Fonti-*Warts* has already acquired a full or even a stronger ability to rescue YAP’s cytoplasmic localization in the LATS1/2 DKO HEK293A cells as compared to its human othologous protein LATS1 (Figure 2E). Collectively, these results not only demonstrated the intrinsically conserved Hippo pathway activity for these three Hippo ancestor components, but also indicated a different alteration of their activities in evolution.

### Opposite evolutionary trends exist among the three Hippo pathway ancestor components

Next, we further assessed the role of Fonti-*Warts* by taking human LATS1 as a control in MDA-MB-231, a human triple negative breast cancer cell line with the impaired Hippo signaling activity due to NF2 deficiency(Dupont, et al. 2011) (Figure 3A). Consistent with our reconstitution study (Figure 2E), Fonti-*Warts* showed a stronger ability than human LATS1 in inducing YAP phosphorylation at S127 (Figure 3A), promoting YAP’s cytoplasmic translocation (Figure 3B) and inhibiting YAP downstream gene transcription (Figure 3C). *Warts*/LATS activation requires its association with adaptor *Mats*/MOB1(Wei, et al. 2007; Ni, et al. 2015) and the NF2-mediated its membrane translocation and the followed *Hippo*/MST-induced phosphorylation(Yin, et al. 2013). Interestingly, Fonti-*Warts* interacted with human MOB1 (Figure 3D) and showed membrane localization even in the NF2-deficient MDA-MB-231 cells (Figure 3B), suggesting that Fonti-*Warts* has already acquired the basic properties required for its activation. Moreover, Fonti-*Warts* displayed much higher kinase activity than human LATS1 by comparing their abilities to phosphorylate the bacterially purified GST-YAP (Figure 3E) and induce their autophosphorylation (Figure 3F). These data suggested a decrease of *Warts*/LATS kinase activity from its unicellular ancestor in evolution.

**Figure 3.**
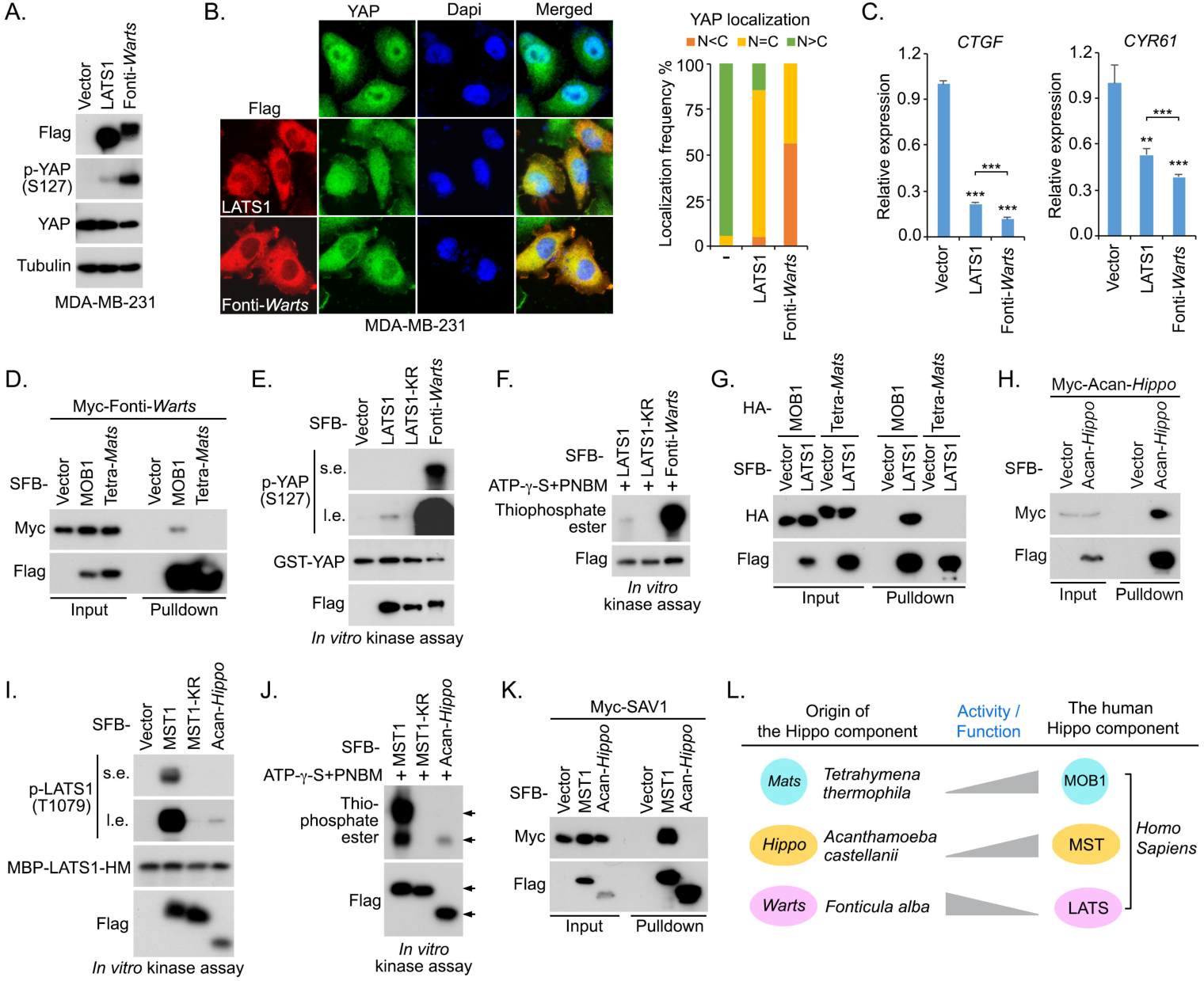
Characterization of the Hippo pathway unicellular ancestor components. (This figure is related to Figures S2 and S3). (**A-C**) *Fonticula alba Warts* (Fonti-*Warts*) has a stronger ability than human LATS1 to phosphorylate YAP at S127 (**A**), translocate YAP into the cytoplasm (**B**) and inhibit YAP downstream gene transcription (**C**) in MDA-MB-231 cells. Flag-positive cells (~200 cells in total) were randomly selected and quantified for YAP localization. ** *p* < 0.01, *** *p* < 0.001 (Student’s *t*-test). (**D**) Fonti-*Warts* interacts with human MOB1. HEK293T cells were transfected with the indicated constructs and subjected to the pulldown assay. (**E** and **F**) Fonti-*Warts* has a stronger kinase activity than human LATS1. SFB-LATS1, SFB-LATS1 kinase-dead mutant (LATS1-KR) and SFB-Fonti-*Warts* were purified from HEK293T cells by using S protein beads and washed thoroughly with high-salt buffer containing 250 mM NaCl. *In vitro* kinase assay was performed to examine their abilities to phosphorylate the bacterially purified GST-YAP (**E**) or induce the auto-phosphorylation (**F**). s.e., short exposure. l.e., long exposure. **(G)** *Tetrahymena thermophile Mats* (Tetra-*Mats*) failed to bind human LATS1. HEK293T cells were transfected with the indicated constructs and subjected to the pulldown assay. **(H)** *Acanthamoeba castellanii Hippo* (Acan-*Hippo*) can form as a dimer. HEK293T cells were transfected with the indicated constructs and subjected to the pulldown assay. (**I** and **J**) Acan-*Hippo* has a weaker kinase activity than human MST1. SFB-MST1, SFB-MST1 kinase-dead mutant (MST1-KR) and SFB-Acan-*Hippo* was purified from HEK293T cells by using S protein beads and washed thoroughly with high-salt buffer containing 250 mM NaCl. *In vitro* kinase assay was performed to examine their abilities to phosphorylate the bacterially purified MBP-LATS1 hydrophobic motif (HM) (**I**) or induce the autophosphorylation (**J**). s.e., short exposure. l.e., long exposure. **(K)** Acan-*Hippo* fails to bind human SAV1. HEK293T cells were transfected with the indicated constructs and subjected to the pulldown assay. **(L)** The activities of three Hippo pathway ancestor components were evolved differently to their human orthologs. The intrinsic activities of two Hippo pathway ancestor components Tetra-*Mats* and Acan-*Hippo* are lower than their respective human orthologs MOB1 and MST1, suggesting that they improved/acquired their abilities to regulate the Hippo pathway in evolution. The intrinsic activity of the Hippo pathway ancestor component Fonti-*Warts* is higher than its human ortholog LATS1, suggesting that its activity was faded to regulate the Hippo pathway in evolution.

Since *Mats*/MOB1 and *Hippo*/MST are two key regulators for *Warts*/LATS, we also characterized their unicellular ancestor proteins Tetra-*Mats* and Acan-*Hippo* by taking their human orthologs MOB1 and MST1 as controls, respectively. In contrast to human MOB1, we hardly detected the interaction between Tetra-*Mats* and human LATS1 (Figure 3G), which may explain why Tetra-*Mats* only partially rescued YAP’ cytoplasmic localization in the MOB1A/B DKO cells (Figure 2C). Moreover, Tetra-*Mats* failed to bind Fonti-*Warts* (Figure 3D). On the other hand, the activation of *Hippo*/MST is regulated by its dimerization and autophosphorylation(Creasy, et al. 1996; Praskova, et al. 2004; Jin, et al. 2012; Deng, et al. 2013; Ni, et al. 2013). Although Acan-*Hippo* can form as a dimer (Figure 3H), its intrinsic kinase activity is relatively low as compared with human MST1 (Figures 3I and 3J). Moreover, Acan-*Hippo* failed to form a complex with human adaptor protein SAV1 (Figure 3K). These functional deficiencies could explain why Acan-*Hippo* has limited ability to rescue YAP’s cytoplasmic localization in the MST/MAP4K-8KO cells (Figure 2D). These data indicated that both *Mats*/MOB1 and *Hippo*/MST may gradually acquire the abilities to regulate *Warts*/LATS from their unicellular ancestors in evolution.

Collectively, our findings revealed opposite evolutionary directions for the three Hippo pathway ancestor components. As for both Tetra-*Mats* and Acan-*Hippo*, their intrinsic activities/functions were further acquired or strengthened when evolved to their human orthologs (Figure 3L); while the intrinsic activity of Fonti-*Warts* was diluted in this evolutionary process (Figure 3L). Given the crucial role of *Warts*/LATS in relaying the upstream signaling to downstream gene transcription in the Hippo pathway, these opposite evolutionary trends may pioneer the fine-tuned regulation of LATS kinase in human.

### Exploring the oncogenic alteration of the Hippo pathway in evolution

Accumulating evidences suggest that dysregulation of the Hippo pathway and hyperactivation of YAP have been frequently associated with various types of human cancers including breast, liver, lung, ovary, colon, kidney, brain and others(Yu, et al. 2015; Zanconato, et al. 2016; Zheng and Pan 2019). Moreover, recent human cancer genome studies also connected the genetic alteration of the Hippo pathway components with cancer development(Sanchez-Vega, et al. 2018; Wang, et al. 2018). Given the evolutionarily conserved roles for the Hippo pathway components, it will be interesting to characterize the oncogenic alteration of the Hippo pathway in evolution.

To achieve this, we profiled the protein sequences of the Hippo pathway core components from unicellular species to human, assigned each amino acid residue in these components with a conservation score, and analyzed the conservative distribution of the Hippo pathway missense mutations derived from two cancer genome databases TCGA and COSMIC (Figure 4A). Actually, these cancer-derived missense mutations were widely distributed through the whole protein sequences, but not significantly correlated with the conservation of the mutated residues (Figure 4A and 4B). These results suggested that the Hippo pathway missense mutations could be randomly occurred in tumorigenesis. Interestingly, when all the TCGA missense mutations were experimentally characterized using a series of Hippo pathway component KO cells(Han, et al. 2018), a group of missense mutations that can functionally inhibit the Hippo signaling activity were found highly enriched in the residues with high conservation scores (Figure 4B). These findings highlighted the significant role of evolutionarily conserved residues in maintaining a normal tumor suppressive function of the Hippo pathway.

**Figure 4.**
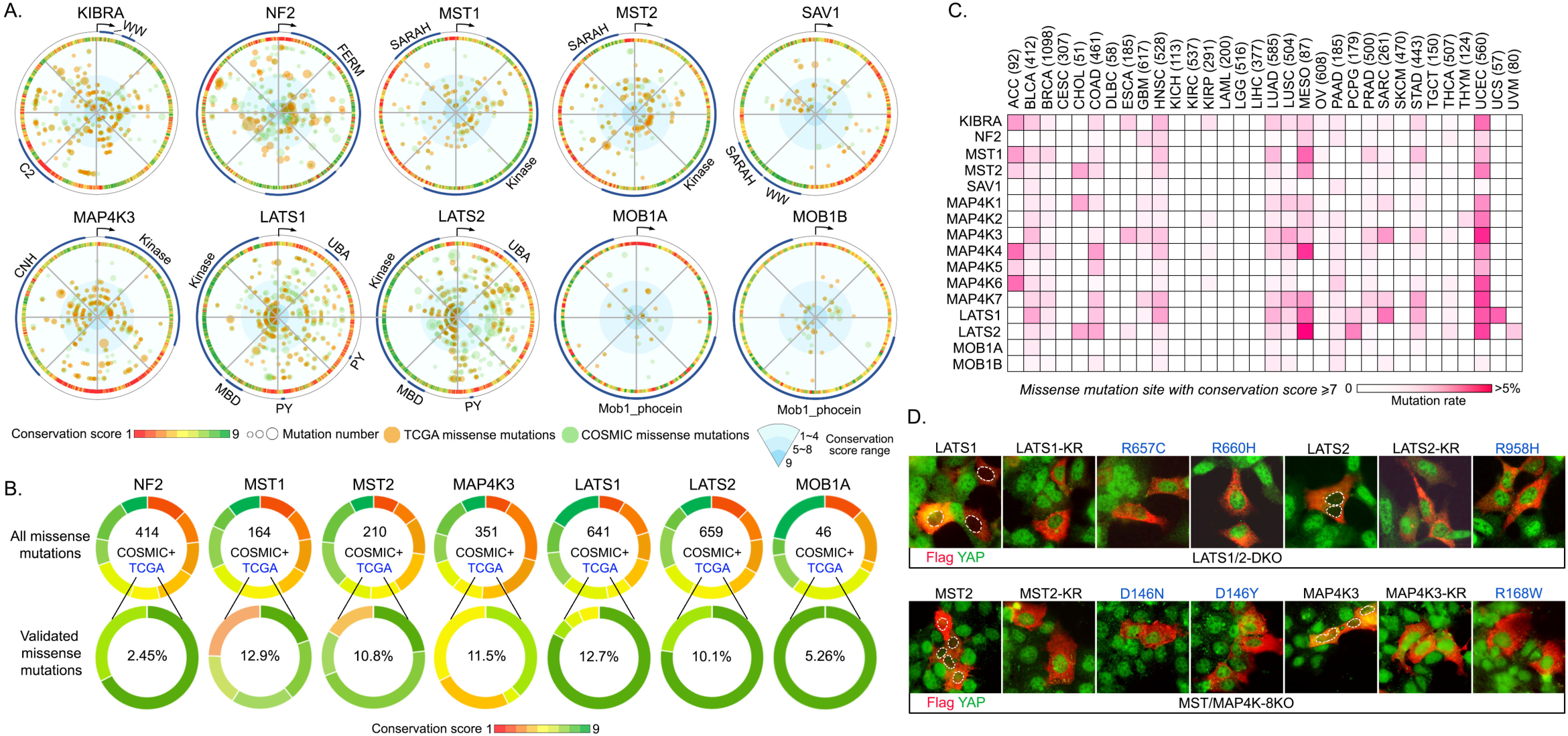
Evolutionary analysis of the Hippo pathway in the human cancer genome. (This figure is related to Table S5) **(A)** A summary of the cancer-derived missense mutations and amino acid conservation information for the indicated Hippo pathway components. The cancer-derived missense mutation data were downloaded from TCGA (labeled in orange) and COSMIC (labeled in green). The dot size is proportional to the mutation number. The conservation scores (1~9; score 1 represents the most variable sites and score 9 represents the most conserved sites in evolution) were obtained in Consurf server and labeled in different colors. The key protein domains and the start amino acid (arrow) were indicated in the outer ring. **(B)** The Hippo signaling-associated functional missense mutations are enriched in the evolutionarily conserved sites. The distribution of the cancer-derived missense mutations was illustrated based on the site conservation scores for the indicated Hippo pathway component. The experimentally validated TCGA missense mutations were enriched in the sites with high conservation scores. The number of total missense mutations in TCGA and COSMIC and the ratio of the experimentally validated TCGA missense mutations were shown. **(C)** Analysis of the evolutionarily conserved missense mutation sites for the Hippo pathway components across 32 TCGA human cancer types. The total case number for each indicated cancer type is shown. The missense mutation sites with their conservation scores at least 7 were included for the analysis. **(D)** Experimental validation of the UCEC-derived missense mutations with high conservation scores in the Hippo pathway component KO cells. The SFB-tagged UCEC-derived missense mutants (with conservation scores ≥7) for LATS1, LATS2, MST1, MST2 and MAP4K3 were expressed in the indicated LATS1/2 DKO and MST/MAP4K-8KO HEK293A cells and labeled in blue.

Next, we subjected the Hippo-associated conserved missense mutations to 32 types of human cancers and found that they were largely enriched in several types of cancers, such as mesothelioma (MESO) and uterine corpus endometrial carcinoma (UCEC) (Figure 4C). MESO is known to contain genetic alteration in multiple Hippo pathway components such as NF2 and LATS2(Sekido 2011). Our comparative analysis further revealed MST1, MAP4K4 and LATS1 as another group of Hippo pathway components, whose conserved missense mutations may also involve in MESO development (Figure 4C). Unexpectedly, UCEC was found significantly enriched with conserved missense mutations in multiple Hippo pathway components including KIBRA, MST1, MST2, MAP4K3, MAP4K6, MAP4K7, LATS1 and LATS2 (Figure 4C). Since Hippo signaling dysregulation has not been well characterized in UCEC, next, we experimentally tested some of the conserved missense mutations for several frequently-mutated Hippo pathway components in UCEC. As shown in Figure 4D, these tested missense mutations all failed the Hippo pathway components LATS1, LATS2, MST2 and MAP4K3 to rescue YAP’s cytoplasmic localization in their corresponding KO cells, suggesting that alteration of the Hippo pathway components at their evolutionarily conserved residues could play an important role in UCEC development.

Taken together, our study not only provided a novel insight into the cancer-related Hippo signaling dysregulation from an evolutionary perspective, but also revealed a new pathological relevance for the Hippo pathway in UCEC.

## Discussion

In this study, we focused on the evolutionary history of the Hippo pathway by addressing three questions. First, why is the paralogous gene number for the Hippo pathway components different between *Drosophila* and human? Second, although an intact and conserved Hippo pathway can be traced to unicellular species, how was this pathway evolved to be a functional one from its ancestor components? Third, what is the connection between the Hippo pathway evolution and its oncogenic alteration in cancer development?

As for the current Hippo signaling research, most of our studies are being carried out in the species/model systems ranging from *Drosophila* to mammals. As compared with *Drosophila*, one of the major differences in the mammalian Hippo pathway is the development of paralogous genes for many Hippo pathway components. Our current study revealed that the turning point of this paralogous gene number change happened in the process of whole genome duplication in fish (Figure 1A). In addition to the Hippo pathway, our unpublished data also showed that the increase of paralogous gene number in other key signaling pathways including PI3K-AKT, WNT, NOTCH, TGF-β, was also initiated in fish. Through it, the complexity and stability of these key signaling pathways were further developed, which maximized their abilities to control tissue/organism homeostasis. In addition, we found that TAZ was originated from *Yorkie* during the whole genome duplication in fish (Figures 1D and S1B), providing an evolutionary insight into their redundant roles in the mammalian Hippo pathway.

To trace the origin of the Hippo pathway, we majorly focused on the firstly emerged Hippo ancestor components *Mats*, *Hippo* and *Warts* in the unicellular species *Tetrahymena thermophile*, *Acanthamoeba castellanii* and *Fonticula alba*, respectively (Figure 2A). In contrast to *Capsapsora owczarzaki* that has already developed an intact Hippo pathway(Sebe-Pedros, et al. 2012), these unicellular species only have one or two Hippo pathway components (Figure 2A), allowing us to characterize their intrinsic activities at the “true origin” of the Hippo pathway. Interestingly, these three Hippo pathway ancestor components all showed conserved functions even in human cells (Figure 2B) and shared key domains and regulatory resides with their orthologs in *Capsapsora owczarzaki*, *Drosophila* and human (Figures S2 and S3). These data suggest that the conserved role of the Hippo pathway could be originated from its ancestor components.

As compared to their human orthologs, both *Mats* and *Hippo* unicellular ancestors were evolved in a “forward” direction where their activities in the Hippo pathway were turned to be strengthened in evolution (Figure 3L). Surprisingly, the evolution of *Warts* unicellular ancestor seems to be occurred in a “backward” manner (Figure 3L), since its intrinsic kinase activity was much higher than that of its human ortholog LATS (Figures 3A-3F). These findings implicated that regulation of the *Warts* activity could be a central event driving the Hippo pathway evolution. During the course that *Warts* activity was decreased from its ancestor, the adaptor function of *Mats* and kinase activity of *Hippo* were both strengthened, a process accompanied by the emergence of additional upstream regulators and downstream effectors in the Hippo pathway (Figure 2A). Based on this hypothesis, this activity adjustment process from the three Hippo ancestor components could be gradually compromised in the species where most of the Hippo pathway components have been emerged, such as *Capsapsora owczarzaki*. Indeed, *Capsapsora owczarzaki Hippo* protein has shown a similar level of activity to its *Drosophila* ortholog in regulating *Drosophila Yorkie*(Sebe-Pedros, et al. 2012). Taken together, our findings provide a novel insight into the Hippo pathway evolution from its ancestor components to an intact pathway in unicellular organisms.

Cancer can be considered as part of an evolution that is now taking place in human, where its associated mutations may result in novel functions and selective advantages(Casas-Selves and Degregori 2011; Lacina, et al. 2019). As a well-established tumor suppressor pathway in control of tissue homeostasis, the Hippo pathway kindly plays a negative role in this evolutionary progress. Although the functionally validated missense mutations in the Hippo pathway were highly enriched in the conserved sites (Figure 4B), the ratio is relatively low (Figure 4B) and the associated cancer type number is also limited (Figure 4C). In contrast, a large portion of the cancer-derived missense mutations were found at the sites with low conservation scores (less than 7), but they did not affect the Hippo pathway activity (Figures 4A and 4B). These findings suggest that mutations at the sites with low conservation may result in novel oncogenic functions for the Hippo pathway components to bypass an active Hippo signaling pathway in tumorigenesis.

Here, we systematically analyzed the Hippo pathway in evolution by elucidating its paralogous gene history, tracing its conserved functions to its unicellular ancestor components and evaluating its cancer-related dysregulation from an evolutionary perspective. However, exactly how the Hippo pathway is able to largely reserve its conserved functions from its unicellular ancestor species to human is still a mystery. In addition, as a key signaling pathway in maintaining tissue homeostasis and preventing cancer development, it is possible that the Hippo pathway could also play a role in regulating an organism’s evolution, which deserves further investigation.

## Materials and Methods

### Searches of the Hippo pathway genes in different species

The genome data of unicellular eukaryotes were downloaded from the National Center for Biotechnology Information (https://www.ncbi.nlm.nih.gov/) genome database. For each main invertebrate phylum and vertebrate class, two species were chosen as representative species from unicellular organisms to multicellular organism. Genes encoding the indicated Hippo pathway core components were isolated from genome sequences of the selected species. The protein sequences of the human and mouse Hippo pathway components were used as queries for the TBLASTN search. The blast results were filtered with e-value < 10^−10^. The introns of the predicted sequences were removed by Genewise (https://www.ebi.ac.uk/Tools/psa/genewise/) by taking their human orthologous genes as queries(Birney, et al. 2004). The following unique domains that were confirmed by PFAM and SMART databases(Schultz, et al. 1998; Finn, et al. 2014) were subjected to the gene searches: the TED domain was used to identify *Scalloped*/TEAD; the Mob1 phocein region was used to identify *Mats*/MOB1; both the catalytic domain of the serine/threonine kinase and the MOB1-binding domain were used to identify *Warts*/LATS; both the catalytic domain of the serine/threonine kinases and *C*-terminal SARAH domain were used to identify *Hippo*/MST; both the WW domain and C2 domain were used to identify *Kibra*/WWC; both the catalytic domain of the serine/threonine kinase and CNH domain were used to identify *Misshapen* or *Happyhour*/MAP4K; the B41 and ERM domains were used to identify *Merlin*/NF2; two WW domains and coiled-coil domain were used to identify *Yorkie*/YAP; one WW domain and coiled-coil domain were used to identify TAZ; two WW domains were used to identify *Salvador*/SAV1. All the Hippo pathway candidate genes were further verified via the phylogenetic reconstruction by taking their human genes as controls. Phylogenetic reconstruction for each dataset was conducted as Maximum Likelihood trees using MEGA X with a Tamura-Nei model and 1000 times bootstrap consensus(Kumar, et al. 2018).

### Sequence alignment and phylogenetic analysis

Sequence alignments were performed by ClustalW with default parameters and visualized by BioEdit (http://www.mbio.ncsu.edu/BioEdit/bioedit.html). Homologous domains, motifs and important residues were checked by alignment. The nucleotide sequences were generated based on the aligned protein sequences and used to reconstruct phylogenetic trees. Specifically, the gaps and amino acids with missing information were deleted, allowing all amino acid sequences in a good alignment for phylogenetic reconstruction.

The conservation of the Hippo pathway components was estimated by the maximum likelihood paradigm in ConSurf server(Ashkenazy, et al. 2016) using the multiple sequence alignment (MSA) data from all the selected species. The evolutionary tree for each dataset was generated based on the species’ evolutionary relationship as estimated by Timetree (http://www.timetree.org)(Kumar, et al. 2017). The conservation score is a relative value and calculated by ConSurf. Based on the relative degree of conservation, each amino acid site was assigned with a value from 1 to 9. Score one indicated the most variable sites while score 9 represented the most conserved sites. Functional divergence between YAP and TAZ was estimated by the program DIVERGE 3.0(Gu, et al. 2013).

### Characterization of the genes saved after whole genome duplication

We used the Ohnologs(Singh, et al. 2015), a database of paralogous genes that were remained from whole genome duplication, to calculate the q score between the Hippo pathway genes and their homologous genes, and quantitatively assess the statistical significance of each paralogous pair. To identify syntenic clusters among the *YAP* and *TAZ* genomic regions in the indicated species, we used BioMart(Smedley, et al. 2009) to get the information of around 1000 genes surrounding the genes of interest and matched them based on the homolog gene information obtained from BioMart. The syntenic clusters were visualized using R packages genoPlotR and OmicCircos(Guy, et al. 2010; Hu, et al. 2014).

### Coevolution analysis

Genetic distance from the multiple sequence alignment data was generated by Mega X with default parameters(Kumar, et al. 2018). To examine the co-evolution of the Hippo pathway core components, we used linear regression analysis to determine the correlation between pairwise evolutionary distances among the Hippo pathway components. Pearson’s correlation coefficient (r) was generated as the output of linear regression analysis to assess the overall evolutionary correlation between any two-gene pairs of interest.

### Antibodies and chemicals

For Western blotting, anti-α-tubulin (T6199, 1:5000 dilution) and anti-Flag (M2) (F3165-5MG, 1:5000 dilution) antibodies were obtained from Sigma-Aldrich. Anti-Myc (sc-40, 1:500 dilution) and anti-GST antibodies (sc-138, 1:1000 dilution) were purchased from Santa Cruz Biotechnology. Anti-phospho-YAP (S127) (4911S, 1:1000 dilution) and anti-phospho-LATS1 (Thr1079) (8654S, 1:1000 dilution) antibodies were purchased from Cell Signaling Technology. Anti-hemagglutinin (HA) monoclonal antibody (901514, 1:2000 dilution) was obtained from BioLegend. Anti-Thiophosphate ester antibody (ab92670, 1:1000 dilution) was purchased from Abcam. The YAP and MBP polyclonal antibodies were generated as previously described(Wang, et al. 2011; Han, et al. 2018).

For immunostaining, an anti-YAP (sc-101199, 1:200 dilution) monoclonal antibody was purchased from Santa Cruz Biotechnology. Anti-Flag polyclonal antibody (F7425, 1:5000 dilution) was obtained from Sigma-Aldrich.

ATP-γ-S kinase substrate (ab138911) and p-Nitrobenzyl mesylate (ab138910) were obtained from Abcam.

### Constructs and viruses

Plasmids encoding the indicated genes were obtained from the Human ORFeome V5.1 library or purchased from Harvard Plasmid DNA Resource Core. For *Tetrahymena thermophila Mats*, *Acanthamoeba castellanii Hippo* and *Fonticula alba Warts*, their sequences were optimized for expression in *Homo sapiens*, synthesized and cloned into a mammalian gene expression vector. All constructs were generated via polymerase chain reaction (PCR) and subcloned into a pDONOR201 vector using Gateway Technology (Invitrogen) as the entry clones. Gateway-compatible destination vectors with the indicated SFB tag, HA tag, Myc tag, GST tag or MBP tag were used to express various fusion proteins. PCR-mediated mutagenesis was used to generate all the indicated site mutations.

### Cell culture and transfection

HEK293T cells were purchased from ATCC and kindly provided by Dr. Junjie Chen (MD Anderson Cancer Center). HEK293A cells were purchased from ThermoFisher and kindly provided by Dr. Jae-Il Park (MD Anderson Cancer Center). MDA-MB-231 cells were purchased from ATCC and kindly provided by Dr. Mien-Chie Hung (MD Anderson Cancer Center). HEK293T, HEK293A and MDA-MB-231 cells were maintained in Dulbecco’s modified essential medium (DMEM) supplemented with 10% fetal bovine serum at 37°C in 5% CO_2_ (v/v) and 1% penicillin and streptomycin.

Plasmid transfection was performed using polyethylenimine and Lipofectamine 3000 (ThermoFisher).

### Immunofluorescent staining

Immunofluorescent staining was performed as described previously(Wang, et al. 2008). Briefly, cells cultured on coverslips were fixed with 4% paraformaldehyde for 10 minutes at room temperature and then extracted with 0.5% Triton X-100 solution for 5 minutes. After blocking with Tris-buffered saline with Tween 20 containing 1% bovine serum albumin, the cells were incubated with primary antibodies for 1 hour at room temperature. After that, the cells were washed and incubated with fluorescein isothiocyanate- and rhodamine-conjugated secondary antibodies for 1 hour. To visualize nuclear DNA, cells were counterstained with 100 ng/mL 4′,6-diamidino-2-phenylindole (DAPI) for 2 minutes. The coverslips were mounted onto glass slides with an anti-fade solution and visualized under a Nikon Ti2-E inverted microscope.

### Gene inactivation by CRISPR/Cas9 system

The indicated MOB1A/B and LATS1/2 double knockout HEK293A cells were generated through the CRISPR/Cas9 system as described previously(Han, et al. 2018). Briefly, five distinct single-guide RNAs (sgRNA) were designed by CHOPCHOP website (https://chopchop.rc.fas.harvard.edu), cloned into lentiGuide-Puro vector (Addgene plasmid # 52963), and transfected into HEK293A cells with lentiCas9-Blast construct (Addgene plasmid # 52962). The next day, cells were selected with puromycin (2 μg/ml) for two days and subcloned to form single colonies. Knockout cell clones were screened by Western blotting to verify the loss of targeted protein expression. The MST/MAP4K-8KO HEK293A cells were kindly provided by Dr. Kun-Liang Guan (University of California at San Diego)

### RNA extraction, reverse transcription and real-time PCR

RNA samples were extracted with TRIzol reagent (Invitrogen). Reverse transcription assay was performed using the Script Reverse Transcription Supermix Kit (Bio-Rad) according to the manufacturer’s instructions. Real-time PCR was performed using Power SYBR Green PCR master mix (Applied Biosystems). For quantification of gene expression, the 2^−ΔΔCt^ method was used. *GAPDH* expression was used for normalization.

The sequence information for each primer used for gene expression analysis is as follows:

*CTGF*-Forward: 5′-CCAATGACAACGCCTCCTG-3′;

*CTGF*-Reverse: 5′-GAGCTTTCTGGCTGCACCA-3′;

*CYR61*-Forward: 5′-AGCCTCGCATCCTATACAACC-3′;

*CYR61*-Reverse: 5′-GAGTGCCGCCTTGTGAAAGAA-3′;

*GAPDH*-Forward: 5′-ATGGGGAAGGTGAAGGTCG-3′;

*GAPDH*-Reverse: 5′-GGGGTCATTGATGGCAACAATA-3′.

### *In vitro* kinase assay

SFB-LATS1 or its kinase-dead mutant (K734R), SFB-MST1 or its kinase-dead mutant (K59R), SFB-*Acanthamoeba castellanii Hippo* and SFB-*Fonticula alba Warts* were expressed in HEK293T cells for 48 hours. The indicated kinases were then pulled down by S protein beads, washed three times in washing buffer (40 mM HEPES, 250 mM NaCl), once in kinase buffer (30 mM HEPES, 50 mM potassium acetate, 5 mM MgCl2), and subjected to the kinase assay in the presence of cold ATP (500 μM) and 2 μg bacterially-purified GST-YAP or MBP-LATS1-C3 (a region containing LATS1 hydrophobic motif)(Han, et al. 2018). The reaction mixture was incubated at 30 °C for 30 min, terminated with 2x SDS loading buffer and subjected to SDS– PAGE. Phosphorylation of YAP and LATS1-HM were determined by YAP S127 and LATS1 T1079 phospho-antibodies, respectively. As for ATP-γ-S-based kinase assay, the purified SFB-tagged kinases were subjected to kinase assay in the presence of 500 μM ATP-γ-S. The reaction mixture was incubated at 30 °C for 30 min, supplemented with 2.5 mM p-Nitrobenzyl mesylate (PNBM) at room temperature for 1h, terminated with 2x SDS loading buffer and subjected to SDS–PAGE. The autophosphorylation of the indicated kinases was detected by an anti-Thiophosphate-ester antibody.

### Human cancer genome analyses

The missense mutation information of the Hippo pathway genes was downloaded from cancer genome databases TCGA (via cBioportal) and COSMIC(Gao, et al. 2013; Forbes, et al. 2016), and compared with the gene conservation data for the Hippo pathway components and our unpublished experimental screen data.

### Quantification and statistical analysis

Each experiment was repeated twice or more unless otherwise noted. There were no samples excluded for the analyses in this study. Data were analyzed by Student’s *t*-test. SD was used for error estimation. A *p* value < 0.05 was considered statistically significant.

## Author contributions

W.W. conceived and designed the study. Y. C. performed the evolutionary analyses. H.H. and W.W. performed the experiments with assistance from G.S., R.V, B.Y. and K.C. H.Z. analyzed the data and revised the manuscript. Y.C and W.W. wrote the manuscript.

## Acknowledgments

We thank Drs. Feng Qiao (University of California, Irvine) and Bao-Liang Song (Wuhan University) for coordinating the Wuhan University visiting student program. This work was supported by a NIH grant (GM126048) and an American Cancer Society Research Scholar Grant (RSG-18-009-01-CCG) to W.W.

---

**Figure S1.**
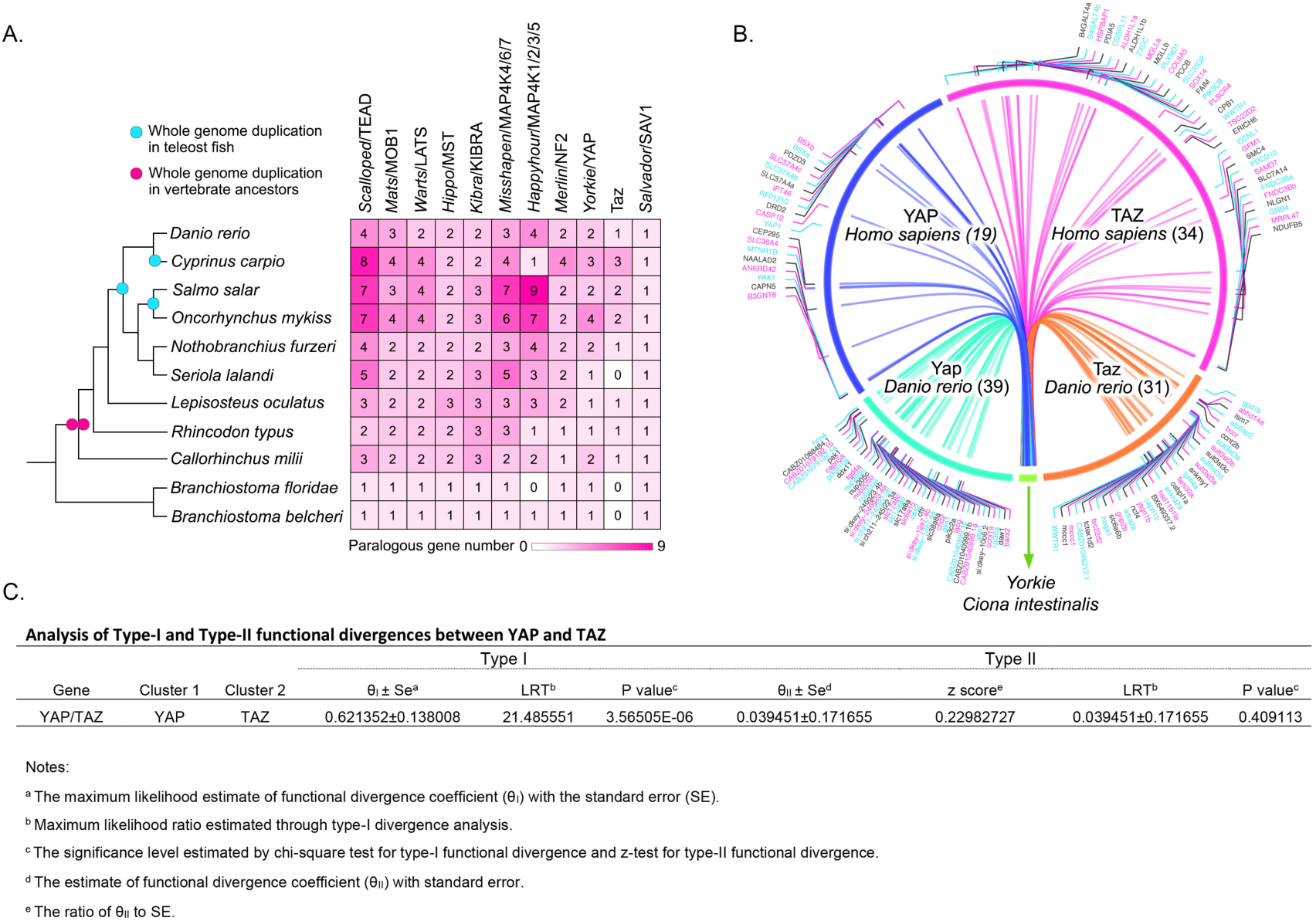
Analysis of the Hippo pathway paralogous gene history in fish. (This figure is related to Figure 1 and Table S1). **(A)** The paralogous gene number of the Hippo pathway components is increased based on the rounds of whole genome duplication in fish. The Hippo pathway core components were searched in the genomes of *Branchiostomidae* and other indicated fish species by TBLASTN, where the human and mouse protein sequences were included as queries. The evolutionary points when the whole genome duplication occurred in teleost fish and vertebrate ancestors were marked as a blue dot and a red dot, respectively. **(B)** The orthologous genes near *YAP* are matched with the ortholog genes near *TAZ* as compared between the genomes from *Homo sapiens* and *Danio rerio*. The syntenic analysis was performed by taking the regions near *Yorkie* in the *Ciona intestinalis* genome as an outgroup. **(C)** The functional divergence test between YAP and TAZ. Both *p* value and parameters used in this test are listed as a table.

**Figure S2.**
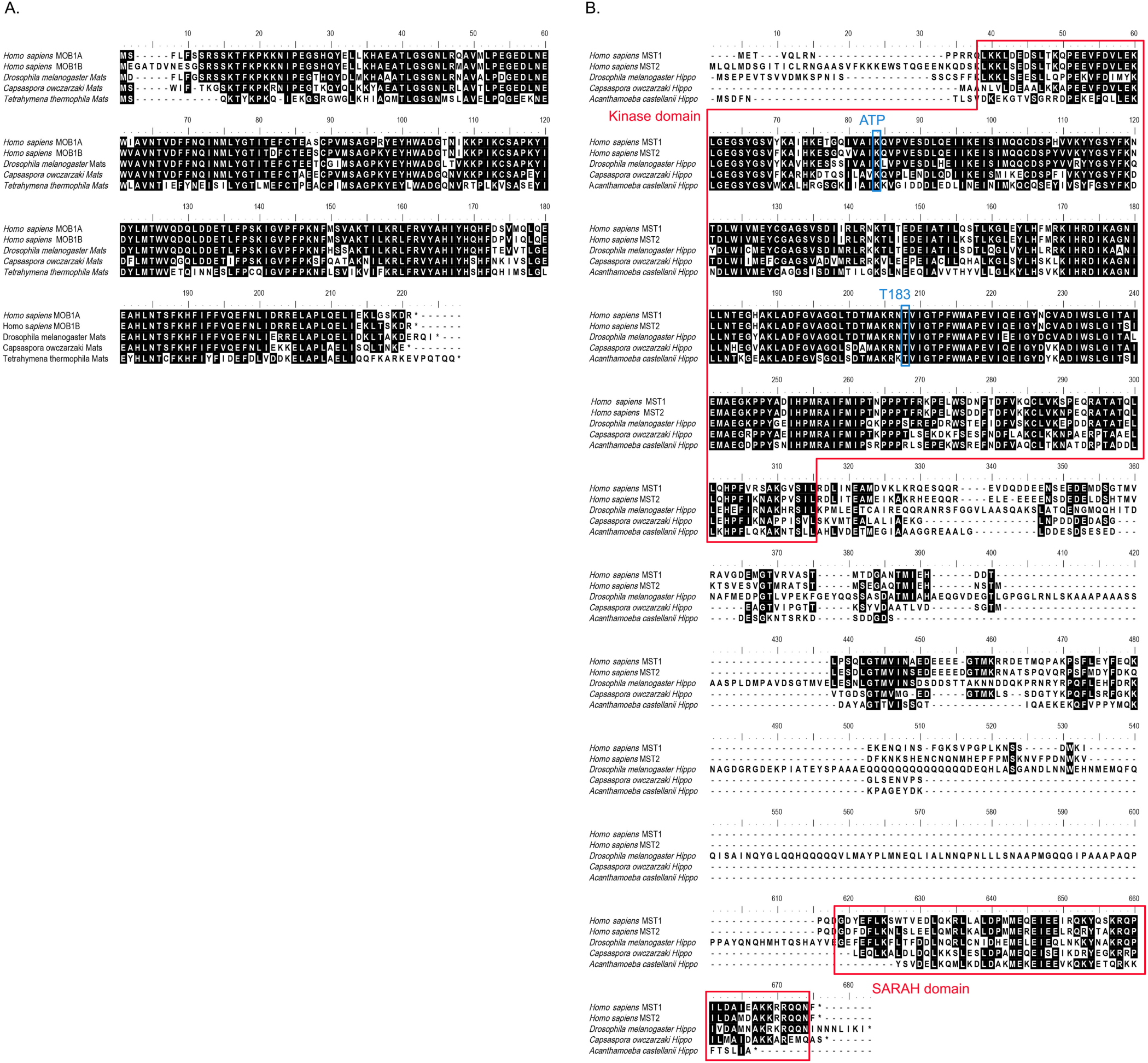
Sequence alignments for *Mats*/MOB1 and *Hippo*/MST proteins from the indicated species. (This figure is related to Figures 2 and 3). **(A)** Sequence alignment for *Mats*/MOB1 is performed for its orthologous proteins in *Homo sapiens*, *Drosophila*, *Capsaspora owczarzaki* and *Tetrahymena thermophile*. **(B)** Sequence alignment for *Hippo*/MST is performed for its orthologous proteins from *Homo sapiens*, *Drosophila*, *Capsaspora owczarzaki* and *Acanthamoeba castellanii*. The *Hippo*/MST kinase domain and SARAH domain were indicated in red. The *Hippo*/MST kinase ATP binding site and autophosphorylation site were indicated in blue.

**Figure S3.**
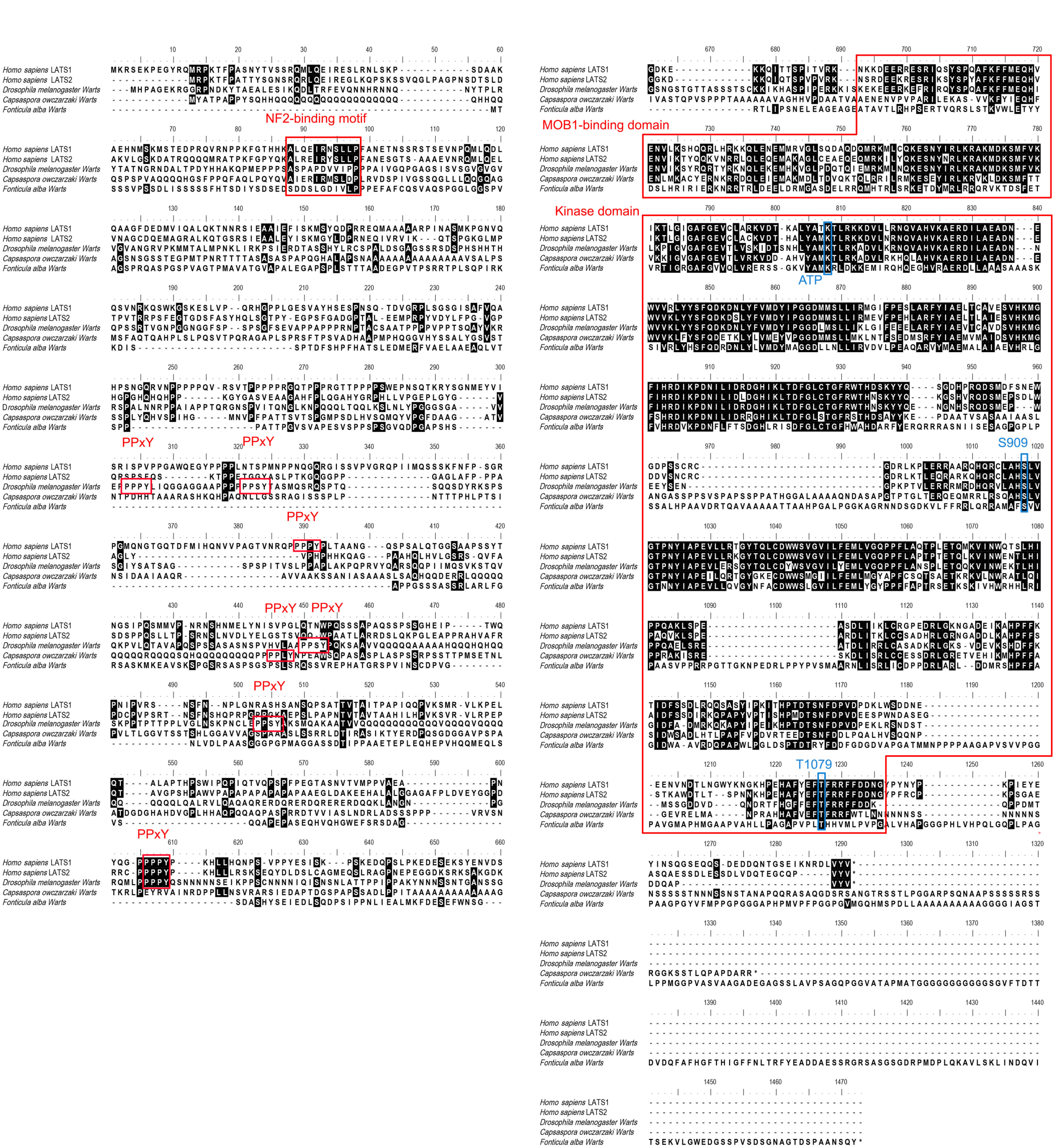
Sequence alignment for *Warts*/LATS proteins from the indicated species. (This figure is related to Figures 2 and 3). Sequence alignment for *Warts*/LATS is performed for its orthologous proteins from *Homo sapiens*, *Drosophila*, *Capsaspora owczarzaki* and *Fonticula alba*. The *Warts*/LATS NF2-binding motif, PPxY motif, MOB1-binding domain and kinase domain were indicated in red. The *Warts*/LATS kinase ATP binding site, autophosphorylation site and phosphorylation site at its hydrophobic motif were indicated in blue.

Table S1. A complete list of DNA sequences used in the phylogenetic analysis. (This table is related to Figures 1, 2, and S1-S3).

Table S2. A complete list of q scores. (This table is related to Figure 1).

Table S3. Pairs of homologous genes near YAP/TAZ/*Yorkie*. (This table is related to Figures 1 and S1).

Table S4. Pairwise genetic distances between ortholog genes in different species and the list of correlation coefficient of coevolution. (This table is related to Figure 2).

Table S5. A complete list of cancer-derived missense mutations and the site conservation scores for the Hippo pathway components. (This table is related to Figure 4).

## References

Ashkenazy H, Abadi S, Martz E, Chay O, Mayrose I, Pupko T, Ben-Tal N. 2016. ConSurf 2016: an improved methodology to estimate and visualize evolutionary conservation in macromolecules. Nucleic Acids Res 44:W344–350.

Birney E, Clamp M, Durbin R. 2004. GeneWise and Genomewise. Genome Res 14:988–995.

Casas-Selves M, Degregori J. 2011. How cancer shapes evolution, and how evolution shapes cancer. Evolution (N Y) 4:624–634.

Conlon I, Raff M. 1999. Size control in animal development. Cell 96:235–244.

Coste A, Jager M, Chambon JP, Manuel M. 2016. Comparative study of Hippo pathway genes in cellular conveyor belts of a ctenophore and a cnidarian. Evodevo 7:4.

Creasy CL, Ambrose DM, Chernoff J. 1996. The Ste20-like protein kinase, Mst1, dimerizes and contains an inhibitory domain. J Biol Chem 271:21049–21053.

Deng Y, Matsui Y, Zhang Y, Lai ZC. 2013. Hippo activation through homodimerization and membrane association for growth inhibition and organ size control. Dev Biol 375:152–159.

Du Q, Kawabe Y, Schilde C, Chen ZH, Schaap P. 2015. The Evolution of Aggregative Multicellularity and Cell-Cell Communication in the Dictyostelia. J Mol Biol 427:3722–3733.

Dupont S, Morsut L, Aragona M, Enzo E, Giulitti S, Cordenonsi M, Zanconato F, Le Digabel J, Forcato M, Bicciato S, et al. 2011. Role of YAP/TAZ in mechanotransduction. Nature 474:179–183.

Finn RD, Bateman A, Clements J, Coggill P, Eberhardt RY, Eddy SR, Heger A, Hetherington K, Holm L, Mistry J, et al. 2014. Pfam: the protein families database. Nucleic Acids Res 42:D222–230.

Forbes SA, Beare D, Bindal N, Bamford S, Ward S, Cole CG, Jia M, Kok C, Boutselakis H, De T, et al. 2016. COSMIC: High-Resolution Cancer Genetics Using the Catalogue of Somatic Mutations in Cancer. Curr Protoc Hum Genet 91:10 11 11–10 11 37.

Gao J, Aksoy BA, Dogrusoz U, Dresdner G, Gross B, Sumer SO, Sun Y, Jacobsen A, Sinha R, Larsson E, et al. 2013. Integrative analysis of complex cancer genomics and clinical profiles using the cBioPortal. Sci Signal 6:pl1.

Glasauer SM, Neuhauss SC. 2014. Whole-genome duplication in teleost fishes and its evolutionary consequences. Mol Genet Genomics 289:1045–1060.

Gu X, Zou Y, Su Z, Huang W, Zhou Z, Arendsee Z, Zeng Y. 2013. An update of DIVERGE software for functional divergence analysis of protein family. Mol Biol Evol 30:1713–1719.

Guy L, Kultima JR, Andersson SG. 2010. genoPlotR: comparative gene and genome visualization in R. Bioinformatics 26:2334–2335.

Halder G, Johnson RL. 2011. Hippo signaling: growth control and beyond. Development 138:9–22.

Han H, Qi R, Zhou JJ, Ta AP, Yang B, Nakaoka HJ, Seo G, Guan KL, Luo R, Wang W. 2018. Regulation of the Hippo Pathway by Phosphatidic Acid-Mediated Lipid-Protein Interaction. Mol Cell 72:328–340 e328.

Harvey K, Tapon N. 2007. The Salvador-Warts-Hippo pathway – an emerging tumour-suppressor network. Nat Rev Cancer 7:182–191.

Hilman D, Gat U. 2011. The evolutionary history of YAP and the hippo/YAP pathway. Mol Biol Evol 28:2403–2417.

Hu Y, Yan C, Hsu CH, Chen QR, Niu K, Komatsoulis GA, Meerzaman D. 2014. OmicCircos: A Simple-to-Use R Package for the Circular Visualization of Multidimensional Omics Data. Cancer Inform 13:13–20.

Jennings HS. 1940. The Beginnings of Social Behavior in Unicellular Organisms. Science 92:539–546.

Jin Y, Dong L, Lu Y, Wu W, Hao Q, Zhou Z, Jiang J, Zhao Y, Zhang L. 2012. Dimerization and cytoplasmic localization regulate Hippo kinase signaling activity in organ size control. J Biol Chem 287:5784–5796.

Kumar S, Stecher G, Li M, Knyaz C, Tamura K. 2018. MEGA X: Molecular Evolutionary Genetics Analysis across Computing Platforms. Mol Biol Evol 35:1547–1549.

Kumar S, Stecher G, Suleski M, Hedges SB. 2017. TimeTree: A Resource for Timelines, Timetrees, and Divergence Times. Mol Biol Evol 34:1812–1819.

Lacina L, Coma M, Dvorankova B, Kodet O, Melegova N, Gal P, Smetana K, Jr. 2019. Evolution of Cancer Progression in the Context of Darwinism. Anticancer Res 39:1–16.

Meng Z, Moroishi T, Mottier-Pavie V, Plouffe SW, Hansen CG, Hong AW, Park HW, Mo JS, Lu W, Lu S, et al. 2015. MAP4K family kinases act in parallel to MST1/2 to activate LATS1/2 in the Hippo pathway. Nat Commun 6:8357.

Ni L, Li S, Yu J, Min J, Brautigam CA, Tomchick DR, Pan D, Luo X. 2013. Structural basis for autoactivation of human Mst2 kinase and its regulation by RASSF5. Structure 21:1757–1768.

Ni L, Zheng Y, Hara M, Pan D, Luo X. 2015. Structural basis for Mob1-dependent activation of the core Mst-Lats kinase cascade in Hippo signaling. Genes Dev 29:1416–1431.

Olson BJ. 2013. Multicellularity: From brief encounters to lifelong unions. Elife 2:e01893.

Pan D. 2010. The hippo signaling pathway in development and cancer. Dev Cell 19:491–505.

Plouffe SW, Lin KC, Moore JL, 3rd, Tan FE, Ma S, Ye Z, Qiu Y, Ren B, Guan KL. 2018. The Hippo pathway effector proteins YAP and TAZ have both distinct and overlapping functions in the cell. J Biol Chem 293:11230–11240.

Praskova M, Khoklatchev A, Ortiz-Vega S, Avruch J. 2004. Regulation of the MST1 kinase by autophosphorylation, by the growth inhibitory proteins, RASSF1 and NORE1, and by Ras. Biochem J 381:453–462.

Sanchez-Vega F, Mina M, Armenia J, Chatila WK, Luna A, La KC, Dimitriadoy S, Liu DL, Kantheti HS, Saghafinia S, et al. 2018. Oncogenic Signaling Pathways in The Cancer Genome Atlas. Cell 173:321–337 e310.

Schultz J, Milpetz F, Bork P, Ponting CP. 1998. SMART, a simple modular architecture research tool: identification of signaling domains. Proc Natl Acad Sci U S A 95:5857–5864.

Sebe-Pedros A, Zheng Y, Ruiz-Trillo I, Pan D. 2012. Premetazoan origin of the hippo signaling pathway. Cell Rep 1:13–20.

Sekido Y. 2011. Inactivation of Merlin in malignant mesothelioma cells and the Hippo signaling cascade dysregulation. Pathol Int 61:331–344.

Singh PP, Arora J, Isambert H. 2015. Identification of Ohnolog Genes Originating from Whole Genome Duplication in Early Vertebrates, Based on Synteny Comparison across Multiple Genomes. PLoS Comput Biol 11:e1004394.

Smedley D, Haider S, Ballester B, Holland R, London D, Thorisson G, Kasprzyk A. 2009. BioMart--biological queries made easy. BMC Genomics 10:22.

Srivastava M, Simakov O, Chapman J, Fahey B, Gauthier ME, Mitros T, Richards GS, Conaco C, Dacre M, Hellsten U, et al. 2010. The Amphimedon queenslandica genome and the evolution of animal complexity. Nature 466:720–726.

Tan L, Nomanbhoy T, Gurbani D, Patricelli M, Hunter J, Geng J, Herhaus L, Zhang J, Pauls E, Ham Y, et al. 2015. Discovery of type II inhibitors of TGFbeta-activated kinase 1 (TAK1) and mitogen-activated protein kinase kinase kinase kinase 2 (MAP4K2). J Med Chem 58:183–196.

Tinti M, Dissanayake K, Synowsky S, Albergante L, MacKintosh C. 2014. Identification of 2R-ohnologue gene families displaying the same mutation-load skew in multiple cancers. Open Biol 4:140029.

Wang W, Chen L, Ding Y, Jin J, Liao K. 2008. Centrosome separation driven by actin-microfilaments during mitosis is mediated by centrosome-associated tyrosine-phosphorylated cortactin. J Cell Sci 121:1334–1343.

Wang W, Huang J, Chen J. 2011. Angiomotin-like proteins associate with and negatively regulate YAP1. J Biol Chem 286:4364–4370.

Wang Y, Xu X, Maglic D, Dill MT, Mojumdar K, Ng PK, Jeong KJ, Tsang YH, Moreno D, Bhavana VH, et al. 2018. Comprehensive Molecular Characterization of the Hippo Signaling Pathway in Cancer. Cell Rep 25:1304–1317 e1305.

Wei X, Shimizu T, Lai ZC. 2007. Mob as tumor suppressor is activated by Hippo kinase for growth inhibition in Drosophila. EMBO J 26:1772–1781.

Yin F, Yu J, Zheng Y, Chen Q, Zhang N, Pan D. 2013. Spatial organization of Hippo signaling at the plasma membrane mediated by the tumor suppressor Merlin/NF2. Cell 154:1342–1355.

Yu FX, Zhao B, Guan KL. 2015. Hippo Pathway in Organ Size Control, Tissue Homeostasis, and Cancer. Cell 163:811–828.

Zanconato F, Cordenonsi M, Piccolo S. 2016. YAP/TAZ at the Roots of Cancer. Cancer Cell 29:783–803.

Zheng Y, Pan D. 2019. The Hippo Signaling Pathway in Development and Disease. Dev Cell 50:264–282.

Zheng Y, Wang W, Liu B, Deng H, Uster E, Pan D. 2015. Identification of Happyhour/MAP4K as Alternative Hpo/Mst-like Kinases in the Hippo Kinase Cascade. Dev Cell 34:642–655.

